# Evaluation of oral baits and distribution methods for Tasmanian devils (*Sarcophilus harrisii*)

**DOI:** 10.1101/2022.04.13.486902

**Authors:** Sean Dempsey, Ruth J. Pye, Amy T. Gilbert, Nicholas M. Fountain-Jones, Jennifer M. Moffat, Sarah Benson-Amram, Timothy J. Smyser, Andrew S. Flies

**Affiliations:** Menzies Institute for Medical Research, College of Health and Medicine, University of Tasmania, Hobart, TAS, 7000, Australia; School of Natural Sciences, College of Science and Engineering, University of Tasmania, Hobart, TAS, 7001, Australia; National Wildlife Research Center, USDA, APHIS, Wildlife Services, Fort Collins, CO, USA; University of British Columbia

**Author notes:** **Correspondence** Andrew S. Flies, PhD, Menzies Institute for Medical Research, College of Health and Medicine, University of Tasmania, Private Bag 23, Hobart TAS 7000.

**Keywords:** transmissible cancer, devil facial tumour disease, oral vaccine, wildlife disease, bait dispenser, feeding behaviour, landscape vaccine distribution, pest control, captive trials, enteric-coated capsule

## Abstract

**Context:** Diseases are increasingly contributing to wildlife population declines. Tasmanian devil (*Sarcophilus harrisii*) populations have locally declined by 82% largely due to the morbidity and mortality associated with two independent transmissible devil facial tumours (DFT1 and DFT2). Toxic baits are often used as a management tool for controlling vertebrate pest populations in Australia, but in other areas of the world oral baits are also used to deliver vaccines or pharmaceuticals to wildlife. Oral vaccine bait products have been distributed for more than 50 years at a landscape scale to protect wildlife from rabies virus and contributed to the elimination of fox rabies from more than ten European nations. An oral bait vaccine to protect against devil facial tumours has been proposed as a management tool to improve the population health, resiliency and fitness of wild Tasmanian devils.

**Aim:** Our goal was to evaluate the potential use of edible baits as vehicles for vaccine delivery to Tasmanian devils.

**Method:** We tested placebo versions of baits that are already used in Australia.

**Key results:** Captive devils consumed all types of placebo baits but exhibited a preference for ruminant- and fish-based baits. Captive devils also consumed inert capsules inserted into placebo baits. Bait fate trials in the field revealed that baits were generally removed within 24 hours. Tasmanian pademelons (*Thylogale billardierii*), brushtail possums (*Trichosurus vulpecula*), and Eastern quolls (*Dasyurus viverrinus*) were the most common nontarget bait competitors at six private properties in southern Tasmania; wild devils removed approximately 5% of ground baits at these sites. We also evaluated an automated bait dispenser, which reduced nontarget uptake and resulted in over 50% of the baits being removed by devils.

**Conclusions:** This study demonstrates that captive and wild devils will accept and consume placebo versions of commercial baits. Bait dispensers or modified baits or baiting strategies are needed to increase bait uptake by devils.

**Implications:** Bait dispensers can be used at a regional scale to deliver baits to Tasmanian devils. These could act as vehicles for preventive or therapeutic vaccines to mitigate the impacts of disease on devil populations.

**Short summary:** This study aimed to test oral baits as potential vaccine delivery vehicles for Tasmanian devils. Captive and wild devils consumed placebo versions of commercial baits used on mainland Australia. Abundant non-target species, such as brushtail possums, Tasmanian pademelons, and eastern quolls consumed most baits in the wild. Implementation of automated bait dispensers increased bait uptake by devils to over 50% at the same regional field sites.

## Introduction

Disease is increasingly a driver of wildlife declines in Australia (Woinarski *et al*. 2019). Disease-induced population declines are most common when new pathogens emerge or non-endemic pathogens are introduced to wildlife populations that have no prior history with the pathogen. Two completely new pathogens were discovered in Tasmanian devils (*Sarcophilus harrisii*) in 1996 and 2014 (Jones *et al*. 2004; Pye *et al*. 2016). Devil facial tumour 1 (DFT1) is a transmissible cancer that was first observed in 1996 (Pearse & Swift, 2006) and has been the primary driver of an estimated 82% decline in regional devil abundance during the last 25 years (Lazenby *et al*. 2018; Cunningham *et al*. 2021). DFT1 infection is nearly always fatal to devils and has spread across more than 90% of Tasmania (Cunningham *et al*. 2021). A second independent devil facial tumour (DFT2) was discovered in 2014 (Pye *et al*. 2016) and to date has been detected only in southern Tasmania.

The island state of Tasmania is the only place where devils still exist in the wild, making this area a critical conservation priority. In addition to disease, other threats to long-term persistence of devil populations include habitat loss, roadkill (Jones 2000; Hobday and Minstrell 2008), domestic animal predation (Holderness-Roddam and McQuillan 2014), and population genetic inbreeding (Lawrence and Wiersma 2019). The decline of the devil population has altered the ecology of Tasmanian ecosystems and has been associated with a greater abundance of invasive feral cats (*Felis catus*) and reduced abundance of native southern brown bandicoots (*Isoodon obesulus*) (Cunningham *et al*. 2020).

A key management strategy for the persistence of devils has involved breeding of disease-free insurance populations both in captivity and wild-living on Maria Island (Thalmann *et al*. 2016). These insurance populations have been used to supplement genetic diversity of wild populations. However, indefinite maintenance of a captive insurance population is costly and introduced devils can also have negative consequences for other native island fauna (e.g birds) (Scoleri *et al*. 2020). Alternative management strategies must be considered and may be necessary to recover wild devil populations.

Development of a vaccine to protect devils or improve resistance from lethal DFT1 or DFT2 infection would be a valuable management tool. Recent work has shown that the devil immune system can recognise and kill DFT1 cells (Pye et al., 2016b, Hamede et al., 2020). Furthermore, priming the immune system of healthy devils with a vaccine followed by immunotherapy after tumours developed was able to induce complete tumour regressions in captive trials (Tovar *et al*. 2017). Previous field vaccine trials were not efficacious in protecting devils from developing tumours, but increased devil immune recognition of tumours (Pye *et al*. 2018; Pye *et al*. 2021). These findings demonstrate that the devil immune system can be stimulated to protect devils against DFT1. However, delivering vaccines to wild devils throughout the rugged Tasmanian landscape presents additional challenges.

Vaccines have been delivered in baits to control wildlife rabies since the late 1970s. An estimated 665,000,000 oral baits were used in Europe alone during 1978-2014 and resulted in the elimination of rabies in red foxes (*Vulpes vulpes*) from western and central Europe (reviewed in Müller *et al*., 2015). Following the success of the rabies oral bait vaccine (OBV) programs, OBVs have been tested for other wildlife diseases including tuberculosis vaccines for wild boars (*Sus scrofa*) (Ballesteros et al., 2011) and European badgers (Palphramand et al., 2017), and sylvatic plague vaccines for prairie dogs (Rocke et al., 2017). Rabies OBV campaigns have demonstrated success despite the challenges of working with a multi-host pathogen and transboundary zoonosis.

As Tasmania is an island state, targeting of devil facial tumours presents a potentially more tractable problem than rabies. This is supported by two independent analyses that recently suggested that an OBV could potentially eliminate DFTD from Tasmania (Lamp *et al*. 2021; Drawert *et al*. 2022). An OBV platform for DFT1 and DFT2 is in development (Flies *et al*. 2020) but the success of an OBV will depend not only on the effectiveness of the vaccine but also the bait distribution system. OBV strategies will depend on the ecology of the target species, characteristics of the bait matrix, production costs, method of bait delivery, bait distribution system, and the environment where baits are distributed. Unfortunately, little is known about bait preferences and bait uptake in Tasmanian wildlife.

The objective of this study was to evaluate the potential use of edible baits as vaccine delivery vehicles for Tasmanian devils. We tested placebo versions of commercially available toxic baits used for control of vertebrate pests on mainland Australia. We offered baits to captive devils to assess bait palatability, quantify feeding behaviours, and determine suitability and acceptance of inert capsules in baits for downstream vaccine encapsulation. We performed bait fate trials in select habitats in southern Tasmania to assess bait uptake by devils and non-target species. Finally, we evaluated an automated bait dispenser to potentially increase specificity of bait uptake to devils. This study advances proof of concept for an OBV distribution strategy for devils and information about bait uptake which may be relevant to vertebrate pest control programs.

## Materials and methods

### Study sites

Trowunna Wildlife Sanctuary (Mole Creek) is a private property located within a eucalypt forest in north-western Tasmania (**Figure 1**). Data collection at this site occurred during late autumn (May) through mid-winter (July) in 2021. Trowunna was visited three times for a total of nine trial days to test bait palatability and feeding behaviours. Studies with captive devils were also performed at the Department of Natural Resources and Environment Tasmania Cressy Research and Development Station in north-eastern Tasmania (**Figure 1**). Study at this site occurred during May through August. The Cressy facility was visited three times for a total of 11 trial days.

**Figure 1.**
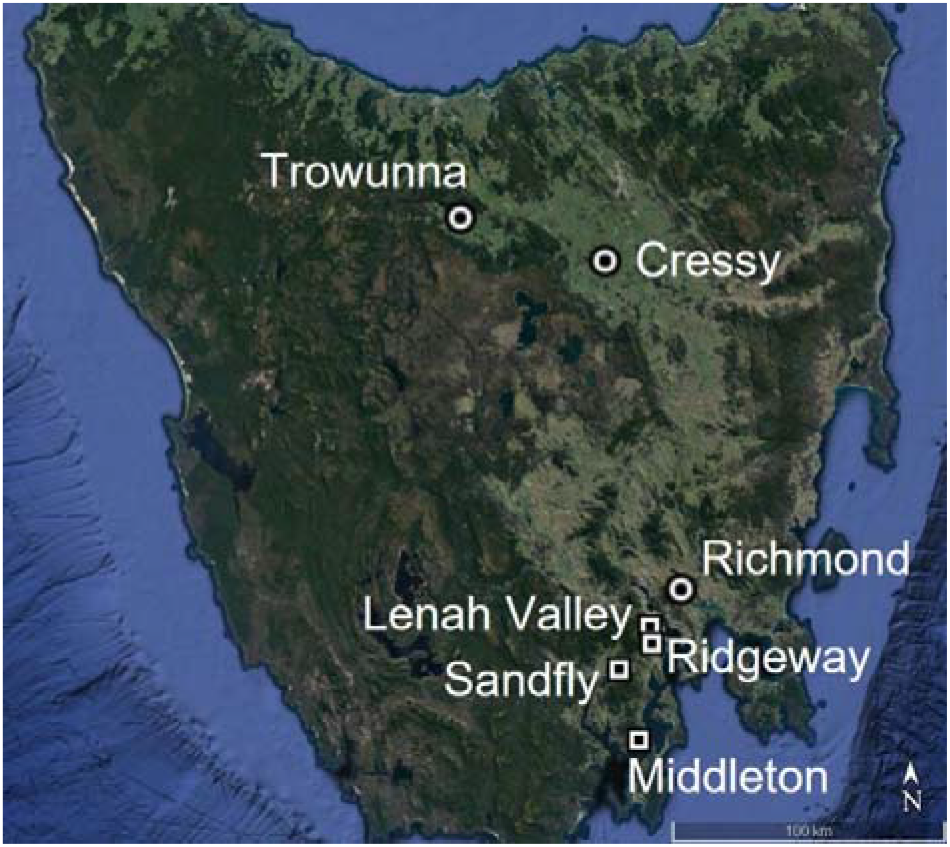
Bait testing locations. Circles indicate relative locations of captive devils. Squares indicates sites used for field testing baits in the wild. Sandfly has three sites and Lenah Valley has two sites. Image created using Google Earth.

All captive devils used for bait trials in this study were individually housed and provided water *ad libitum*. At Trowunna we offered baits without capsules to ten devils and baits containing capsules to five devils. Three of the five devils at Trowunna offered baits without capsules were involved in the initial study without capsules. A minimum period of 60 days elapsed between bait only and bait containing capsule trials. At Cressy we offered baits without capsules to eight devils and baits containing capsules to seven devils. Seven devils were included in both trials with a minimum of 90 days between the two types of trials.

Trowunna devils are typically fed a daily ration consisting primarily of Bennett’s wallaby (*Notamacropus rufogriseus*), Tasmanian pademelon *(Thylogale billardierii)*, brushtail possum (*Trichosurus vulpeca*), rabbit (*Oryctolagus cuniculus*) and a variety of domestic poultry. Devils at Trowunna were fasted 24 hours prior to the first day of each bait trial and were then fed the daily ration at the conclusion of each trial day. Cressy devils are typically fed a diet consisting primarily of brushtail possum, Tasmanian pademelon and Bennett’s wallaby three times per week as well as weekly enrichment feeding such as chicken (*Gallus gallus*) eggs. Cressy devils were fasted 24 hours prior to the first day of each trial and were then offered their food ration according to their usual weekly schedule at the conclusion of each trial day.

Bait uptake by wild devils was tested at six private sites in southern Tasmania (**Figure 1**). Three locations were part of a 150 hectare property near Sandfly that contains three permanent dams, open grassland, and an olive tree grove surrounded by eucalypt forest and at least 500 m from the nearest residence. Individual sites in Lenah Valley, Middleton, and Ridgeway were within 100 m of a residence and were either bordered or surrounded by eucalypt forest. One site in Lenah Valley had equine and sheep paddocks within 100 m and eucalypt forest within 500 m. All sites were selected based upon owner-reported devil sightings on the property prior to this study. Property owners volunteered to place the non-toxic baits and retrieve images from remote cameras at least once per week. One site in Sandfly had a trail camera but was not baited.

### Baits

Animal Control Technologies Australia (Melbourne, Australia) supplied four types of placebo baits for this study (**Figure 2**). These baits have been used in mainland Australia as toxic baits containing sodium monofluoroacetate (known as 1080) for reducing populations of introduced vertebrate pest species (e.g., wild canids, feral pigs). The ruminant-based bait was a rectangular bait weighing approximately 35 g. The fish-based bait was cylindrical and weighed approximately 35 g. The 35 g ruminant and fish baits used for these trials contained green scat marker beads. The cereal-based bait was a fish-flavoured cylindrical bait that weighed 70 g. The cereal baits were cut approximately in half for this study and did not contain scat markers. We also used dried kangaroo baits weighing approximately 40 g. Kangaroo baits were treated as a positive control given that macropod meat is a common prey item in the diet of wild devils. We used only fish-based baits (hereafter referred to as fish baits) for the inert capsule trials. Custom fish baits without scat markers (20 mm height by 30 mm diameter; approximately 17 g) were produced to fit the dimensions of the bait magazine and receptacle of an automated bait dispenser.

**Figure 2.**
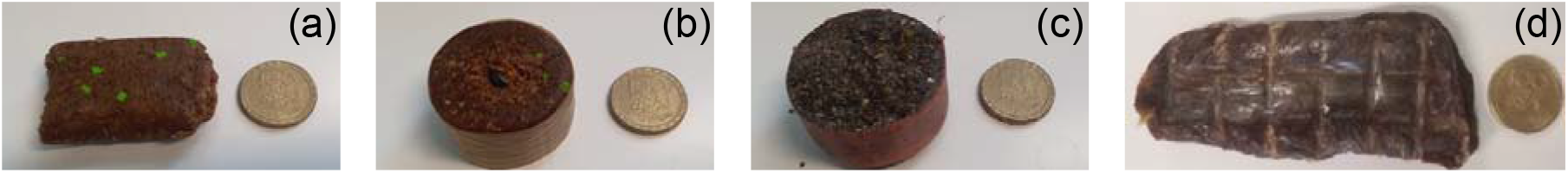
Ruminant meat-based (a), fish-based (b) and fish-flavoured cereal-based (c), and dried kangaroo meat (d) baits. 25 mm coin shown for scale.

### Capsules

We tested inert white-opaque Capsugel Empty DRcaps® capsules made of hypromellose, titanium dioxide and gellan gum (Lonza Australia, #G69CS000753). The enteric coating prevented bait moisture from dissolving the capsules, which otherwise readily dissolve in most liquids. Capsules were filled with a sucrose solution (golden syrup) and coated with fish oil prior to being inserted into a fish bait matrix.

### Video recording in captivity

Cameras were positioned on the boundary of devil pens, which were approximately 1.5 m high, facing the area of bait placement and programmed to record for 30 minutes post-bait offering. Footage was recorded with GoPro8, GoPro9, and DC-FT7 Lumix cameras, and Samsung Note 2, and Samsung S7 phone cameras. Multiple cameras were used to record independent tests simultaneously.

### Bait testing in captivity

Baits were weighed prior to each trial and disposable gloves were worn to avoid adding human scent to the baits. Each captive devil was used in only one trial each day. Handling tools (Nifty Nabber, Unger #92134) were used to simultaneously lower two different bait types into a pen as a two-choice test. Separate tools were used for each bait type to minimise scent transfer. Baits were placed approximately 0.5 m apart. Human presence was minimised during filming but could not be eliminated in every instance (e.g., trials occurring at Trowunna Wildlife Sanctuary occurred during visitation by the public). The behaviours of each devil post bait offering were filmed for approximately 30 minutes.

At the conclusion of filming, pens were checked for presence of baits. Bait remains were weighed to the nearest gram to estimate the percentage consumed. Intact baits and/or bait remains were then returned to the pen to evaluate whether these would be consumed overnight. A bait was recorded as ‘consumed’ only if an individual was observed eating at least part of the bait. The proportion of the bait consumed could not be estimated when the baits were taken off camera and could not be located. These ‘removed’ baits were not included in the results. Bait trials were conducted between 10:00 and 15:00 hours.

In a second set of captive bait trials, a single encapsulated bait was offered in each trial. Fish oil has been reported to be an effective lure, so the capsules were covered in fish oil prior to being inserted into the baits (Andersen *et al*. 2016). We attempted to retrieve the bait and capsule following the trials to weigh baits and determine if the capsule was consumed, punctured, or crushed.

### Video analysis of devil behaviour

Behavioural data were quantified from videos of captive trials using Behavioural Observation Research Interaction Software (BORIS) (Friard and Gamba 2016). All video recordings were imported to BORIS as MP4 files at 30 frames per second. The frequency of ‘point events’ (non-continuous) and the duration (seconds) of ‘state events’ (continuous) were calculated in BORIS. Behaviours were defined based on an ethogram (**Table S1**) developed by Marissa Parrott and staff at the Healesville Sanctuary (Pollock *et al*. 2021). Behaviours not associated with feeding (drinking, bait taken out of frame, devil sniffing the air) were not included in results. The occurrence of behaviours was not mutually exclusive, and up to three events could be logged at the same time.

### Bait testing on private properties

Baits were placed by gloved hand approximately 3 m away from cameras by volunteer landowners on private property. Bait stations were left without baits for a one-week period to ensure cameras were operating adequately. Baits were then placed on the ground with seven-day bait replacement intervals as the first set of treatments. The first two weeks of trials showed baits were mostly consumed or removed by non-target species on the first night, so we modified the trials. For the remaining six weeks, volunteers covered the baits with two cups of soil to limit visual detection of baits as in the prior study that tested ruminant baits in Tasmania (Hughes *et al*. 2011). We also reduced the interval between bait replenishment from seven days to a minimum of 48 hours after disappearance of the prior bait.

A total of 289 nights of footage from six sites were checked for presence of animals. Of these 289 nights, 85 nights had baits present and are referred to as bait exposure nights (BENs). A ‘visit’ was defined as an animal observed on camera at a bait station. This included animals that showed interest in the bait as well as animals that were passing through the site of bait placement. To account for repeat visits, if an individual of the same species was seen multiple times at a bait station within 10 minutes and qualitatively appeared to be of similar size and appearance, it was counted as one visit. Camera traps recorded either 20-second, one-minute, or two-minute videos. Visits by species were taxonomically grouped for analysis as the ‘macropod’ group (Tasmanian pademelon, Bennett’s wallaby, and Tasmanian bettong (*Bettongia gaimardi*)), the ‘other marsupials’ group (southern brown bandicoot (*Isoodon obesulus*) and eastern-barred bandicoot (*Perameles gunnii*)), the ‘rodents’ group (presumable *Rattus* species and *Mus species*), and the ‘bird’ group (Tasmanian native hen (*Tribonyx mortierii*), tawny frogmouth (*Podargus strigoides*) and blackbird (*Turdus merula*)). Brushtail possums and eastern quolls (*Dasyurus viverrinus*) were recorded at the species level due to the relatively high number of visits and baits removed by these species.

An animal was listed as ‘No ID’ when it could not be identified. A bait was marked as ‘suspected taken’ by a particular animal when bait removal could not be confirmed on camera trap footage (e.g. obstructed view), but we had reason to believe that the animal was responsible for the removal (e.g., evidence of small tunnel to the bait suggesting a rodent had removed a bait). When a bait had disappeared from a bait station but there was no footage of the removal, the bait was listed as ‘confirmed removal’ by an unidentifiable animal (No ID). Cameras used to capture footage include one Flag Power camera, four Keep Guard cameras, four Browning BTC-8E cameras, and a set of Arlo Pro 2 cameras.

### Automated bait dispensers

Automated bait dispensers (**Figure S1**) developed for delivering fishmeal polymer baits (e.g., containing oral rabies vaccines) to raccoons (*Procyon lotor*) (Smyser *et al*. 2015) were modified for the delivery of baits to devils. Specifically, the square magazine used for the delivery of cubical baits to raccoons was replaced with 32 mm Vinidex Rural Plus® polyethylene pipe to accommodate the cylindrical 30 mm diameter fish baits used in this study. Ten baits were loaded vertically into each magazine. One of the baits was immediately pushed through to the receptacle platform (and therefore available for consumption) by a motorised piston. A sensor in the receptacle detects when a bait has been removed from the platform and dispenses a new bait 40 minutes after removal of the previous bait. Upon presentation by the dispenser, the bait was available to devils within a polyvinyl chloride (PVC) cylinder (77 mm in diameter × 155 mm in length) to restrict access to non-target species that cannot physically reach the baits with their paw and/or mouth.

Automated bait dispensers were tested at the two Lenah Valley sites and one Sandfly site. Dispensers were secured to posts with rubber straps at either 200 mm or 350 mm above the ground. A camera trap was set up approximately 2-3 m away from the dispenser. Approximately 5 mL fish oil was dispersed around the base of the dispenser to attract devils. Ten custom-made fish baits (20 mm x 30 mm, approximately 17.5 g) were loaded into the dispenser magazine. We monitored bait dispensers with trail cameras for 33 nights. A dispenser interaction was defined as ‘an animal showing interest in and/or contacting the dispenser, including sniffing, making physical contact, or scent marking’.

### Statistical analyses

Data collected from the captive trial data sheets was analysed using R v4.1.1 (R Core Team, 2021). A mixed effects logistic regression model (i.e. a generalised linear mixed model GLMM)) was used to analyse bait preference amongst single devils. The binary response variable was whether a bait was eaten (1) or uneaten (0). Baits were categorised as eaten if more than 50% of the pre-trial weight was consumed. The data were modelled for the response variable using a Bernoulli distribution (log-link). Five models were specified with different combinations of fixed effects. Fixed effects included age, sex, days since the devil’s most recent food ration, bait flavour, site, and temperature. The random effect was the individual animal ID. Smooth splines were fit to model 2 to quantify non-linear effects. We used kangaroo baits as a positive control in only a few trials and all kangaroo baits were consumed; the data from kangaroo baits was not included in models because it was considered impractical to deliver a vaccine within dried meat baits.

We fitted our models using Bayesian inference with Hamiltonian Monte Carlo (HMC) ‘no U-turn sampling’ (NUTS) using the *brms* package (Bürkner 2017) and plotted using *bayesplot* package (Gabry and Mahr 2021). Four Markov chains of 8000 iterations were run using non-informative priors in model specification and every 16^th^ draw was thinned to reduce autocorrelation. Effective sample size (ESS) and R-hat convergence diagnostics revealed no evidence of divergence or autocorrelation in any model (**Figures S2-S4**).

Model selection was performed using approximate leave-one-out (LOO) cross-validation of the posterior predictive log density with the *loo* package (Vehtari *et al*. 2018). The observed values were compared to the modelled predictions using the 95% credibility intervals of the posterior predictive distribution to determine which fixed effects were affecting whether a bait would be eaten in a trial. Model evaluation was validated using area under the curve (AUC) (Delong *et al*. 1988) to measure the ability of the model to correctly distinguish an eaten bait from an uneaten. Missing data were removed before graphing/analysis. Scatterplots illustrating bait preference were made using *ggplot2* package (Wickham 2016).

## Results

### Bait consumption by captive devils

Nineteen of 153 offered baits, including twelve fish, six ruminant, and one kangaroo bait, disappeared from the trial pens, but the animal taking or eating the bait could not be confirmed (**Table S2**). Three cereal baits and one ruminant bait (four total) were partially eaten during trials but crumbled into small pieces that were unable to be weighed and checked for consumption proportions. Circumstantial evidence suggests the baits taken out of view or crumbled were at least partially consumed, but because the baits could not be recovered and weighed they were recorded as ‘removed’ and not included in the analysis. For the baits included in the analysis, complete consumption of baits was observed for 57% (27/46) of ruminant and 53% (20/38) of fish baits compared to 33% (13/40) of the cereal-based baits (**Table S2**). All six kangaroo meat baits were completely consumed. Approximately 15% of the ruminant, fish, and cereal baits were partially consumed (**Table S2**).

The average mass of baits consumed by captive male devils (n=9) was about 25% more than the mass of bait consumed by captive female devils (n=9) for all bait types (**Figure 3**). Amongst male devils, the percent of the bait mass consumed was highest for the fish (73%) and ruminant baits (71%). Ruminant (46%) and fish (39%) baits also had the highest percent consumption by female devils. Interestingly, four-year-old devils (n=3) completely consumed 24/26 baits, whereas six-year-old devils (n=3) consumed only 3/23 baits.

**Figure 3.**
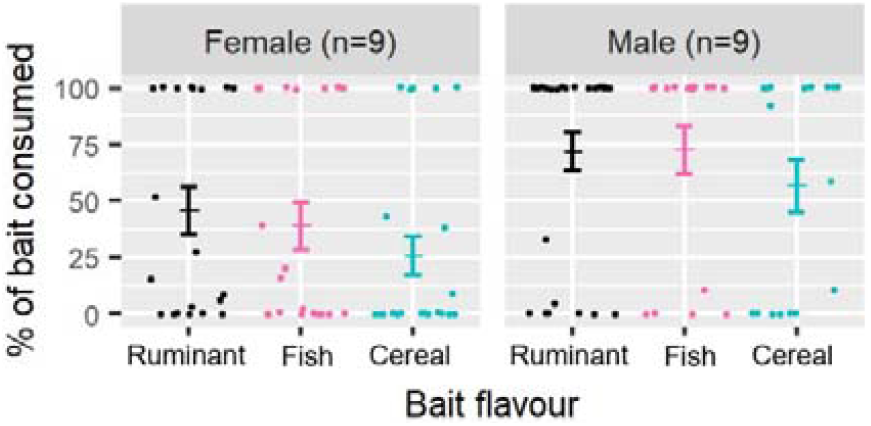
Bait consumption by captive female and male devils. Plots show mean percent consumption by nine female and nine male devils for n=47 ruminant, n=38 fish, and n=40 cereal baits. Dots represent individual bait trials. Vertical bars represent standard error. Horizontal midlines represent mean percent of the bait consumed.

We constructed five models to predict factors associated with whether or not the bait was eaten (**Table 1**). Model performance was measured using expected log pointwise predictive density (ELPD). The results of the leave-one-out cross-validation approach revealed that the simplest linear model (Model 1) had the highest; Model 1 included sex, age, bait flavour and days since fed as fixed effects, along with individual as a random effect. Adding temperature and trial site as fixed effects or using smooth splines for age and days since last fed did not result in any improvement in model performance. Model 1 could predict baits being or uneaten well (area under the curve (AUC) = 0.985, coefficient of determination (Bayesian R2) = 0.76). Captive devils were more likely to eat the ruminant and fish baits than the cereal baits (**Table 2**). There was a moderate indicator for greater percentage consumption by male devils (95% CI -0.84, 30.17) and four-year-old devils (95% credible interval (CI) -15.25, 35.32) (**Table 2**). ‘Days since fed’ was not found to have a strong association with bait uptake.

**Table 1.** Model comparison using leave-one-out (LOO) analysis for 5 models. Model performance was measured using expected log pointwise predictive density (ELPD). Values in the ‘mean (ELPD)’ and ‘std err ELPD’ columns are computed by making pairwise comparisons between each model and the model with the largest.

| Model number | Predictors in model | mean (ELPD) | std err ELPD |
| --- | --- | --- | --- |
| 1 | Sex, Age, Flavour, Days since fed | 0.0 | 0.0 |
| 2 | Sex, Age (spline), Flavour, Days since fed (spline) | -0.2 | 0.9 |
| 3 | Sex, Age, Flavour, Days since fed, Site | -0.8 | 1.1 |
| 4 | Sex, Age, Flavour, Days since fed, Temperature | -2.3 | 1.0 |
| 5 | Sex, Age, Flavour, Days since fed, Temperature, Site | -3.8 | 2.2 |

**Table 2.** Variables are shown with their predicted log odds estimate and 95% credible intervals. Variables whose credible intervals contain zero are considered to have no effect on the predictive performance of the model. See Figure S2 for posterior distributions of each parameter.

| Variable | Log odds estimate | 95% Credible interval |
| --- | --- | --- |
| <u>Fixed effects</u> |  |  |
| Intercept | -1.81 | -23.45, 17.05 |
| Sex: Male | 10.50 | -0.84, 30.17 |
| Age: 2 | -3.46 | -27.11, 19.22 |
| Age: 4 | 8.21 | -15.25, 35.32 |
| Age: 5 | -9.73 | -42.19, 18.87 |
| Age: 6 | -16.42 | -51.56, 7.54 |
| Flavour: fish | -0.23 | -2.32, 1.87 |
| Flavour: cereal | -3.40 | -6.76, -0.96 |
| Days since fed | 1.01 | -0.29, 2.52 |
| <u>Random effects</u> |  |  |
| Individual ID | 8.58 | 3.38, 20.19 |

### Bait and capsule consumption by captive devils

We performed trials with fish baits loaded with an enteric-coated capsule containing golden syrup using 12 singly-housed devils across two sites. We used fish baits for these trials because they performed similarly to ruminant baits in the initial trials, fish oil could be coated on the capsule itself, and the fish baits could be used in subsequent bait dispenser trials more readily than ruminant baits. Fourteen baits were taken out of view and recorded as ‘removed’. In three trials with removed baits, the capsule had separated from the bait and was not consumed. These ‘removed’ baits could not be logged as ‘eaten’, ‘uneaten’, or ‘partially eaten’ and were thus not included in capsule interaction results.

In the 21 capsule trials included in the final analysis, 67% (14/21) of capsule-loaded fish baits were completely eaten, 29% (6/21) were uneaten, and 5% (1/21) were partially eaten. Amongst the 14 eaten baits, 50% (7/14) of capsules were completely consumed, and a further 21% (3/14) were punctured. Capsules separated from the baits in 48% (10/21) of these trials. On the occasions where capsules became separated, 60% (6/10) of capsules were still consumed or punctured.

### Captive devil feeding behaviour

A total of 941 state events were logged from 62 trials in which two baits without capsules were simultaneously offered to 16 individually housed devils. The occurrence of observed behaviours was not mutually exclusive. The ‘bait crushed’ in jaws behaviour comprised the largest proportion of time amongst state events (37%) (**Table 3**). Amongst observable state events (i.e. excluding “bait interaction – obstructed view’), ‘bait held in paws’ was the next most frequently occurring behaviour (19%), followed by ‘bait sniffed’ (12%). Devils displayed a total of seven different point behaviours and 424 point events in bait only trials (**Table 3**). ‘Bait approached’ was the most commonly occurring behaviour (51%). ‘Bait contacted’ also represented a considerable proportion of behaviours (40%). ‘Scratch mouth’, ‘scent mark’, and ‘scratch body’ behaviours made up lesser proportions of point events in both groups.

**Table 3.** Behavioural interactions with baits. Percent of duration shown for state events and percent of events shown for point events.

|  | Bait<br>only | Bait and<br>capsule |
| --- | --- | --- |
| State behaviours | n = 941 | n = 164 |
| bait crushed | 37.2 | 42.0 |
| obstructed view | 23.9 | 13.0 |
| bait held in paws | 18.6 | 26.3 |
| bait sniffed | 11.6 | 2.9 |
| bait nibbled | 4 | 2.1 |
| bait pinned to ground | 1.5 | 0.0 |
| face wipe | 1.1 | 4.5 |
| lick paws | 1.1 | 4.4 |
| bait licked | 0.9 | 1.3 |
| capsule consumed | NA | 0.4 |
| Point behaviours | n = 424 | n = 67 |
| bait approached | 51.4 | 40.3 |
| bait contacted | 39.9 | 37.3 |
| scent mark | 4.0 | 3.0 |
| scratch mouth | 3.1 | 0.0 |
| scratch body | 1.4 | 3.0 |
| enrichment | 0.2 | 0.0 |
| capsule punctured | NA | 16.4 |

A total 164 state behaviours were recorded from 29 capsule bait trials with eleven devils. ‘Capsule contents licked’ is the only behaviour observed in the capsule trials and not in preference trials (**Table 3)**. ‘Bait crushed’ (42%) and ‘bait held in paws’ (26%) again comprised the largest proportion of state behaviours. A total of 67 point events were recorded from capsule bait trials. ‘Bait capsule punctured’ was the only point event observed in capsule trials that was not in the preference trials. ‘Bait approached’ was the most commonly occurring point event (40%) (**Table 3**). ‘Bait contacted’ comprised the next largest proportion of point event (37%). ‘Bait capsule punctured’ comprised 16% of events.

### Wild devil bait trials

Macropods were by far the most common animals observed (60.4%, 822/1361) in videos across 289 nights where bait was present and not present at the six field sites (**Table S3)**; pademelons were the most common macropod, accounting for 56.4% of the total visits (768/1361). Possums were the next most observed animals (11.5%). Eastern quolls and devils made up 7.3% and 4.9% of the total animals observed, respectively.

In the subset of footage from 85 bait exposure nights (BENs) when baits were present, five sites were visited on 325 occasions by individuals from 10 species or higher taxonomic classifications (e.g., macropods). Amongst identifiable animals, 63% (206/325) of animals observed at bait stations when baits were present were macropods and 17%, 6%, and 4% of the remaining visitors on BENs were brushtail possums, devils, and eastern quolls (**Table 4)**. Rabbits, cats, dogs, and other marsupials were observed infrequently, with rodents being observed the least but potentially underestimated due to their smaller size.

**Table 4.** Total bait uptake and visitation. Species/categories of animals that were observed at bait stations on bait exposure nights (BENs) for all bait flavours. Species/category are ranked by number of visits. Interactions are represented as ‘visits/bait taken, with ‘(susp.)’ indicating where a bait is suspected to have been taken by the corresponding species/category of animal.

| Species or group | RUMINANT | FISH | CEREAL | Kangaroo | Total |
| --- | --- | --- | --- | --- | --- |
| BENs | n = 33 | n = 15 | n = 22 | n = 15 | n = 85 |
| Macropods | 86/2 | 26/1 (1) | 88/3 (1) | 6/0 | 206/6 (2) |
| Brushtail possum | 25/2 (1) | 8/2 (1) | 7/1 (1) | 16/0 (1) | 56/5 (4) |
| Tasmanian devil | 2/0 | 4/0 (1) | 12/1 (1) | 2/0 | 20/1 (2) |
| Eastern quoll | 6/2 (2) | 2/2 | 4/2 | 0/0 | 12/6 (2) |
| Birds | 4/0 | 1/0 | 0/0 | 5/0 | 10/0 (0) |
| No ID | 2/4 | 2/4 | 3/4 | 0/4 | 7/16 (0) |
| Rabbits | 4/0 | 0/0 | 0/0 | 0/0 | 4/0 (0) |
| Cat | 1/0 | 0/0 | 2/0 | 0/0 | 3/0 (0) |
| Other marsupials | 0/0 | 0/0 | 3/0 | 0/0 | 3/0 (0) |
| Dog | 1/0 | 1/0 | 1/0 | 0/0 | 3/0 (0) |
| Rodents | 0/0 | 1/0 (1) | 0/0 | 0/0 | 1/0 (1) |
| Total | 131/10 (3) | 45/9 (4) | 120/11 (3) | 29/4 (1) | 325/34 (11) |
| Visits/night | 3.96 | 3 | 5.45 | 1.93 | 3.82 |

Nineteen percent of baits (16/85) were classified as taken by ‘No ID’ because they were removed from the bait station but the animal responsible could not be identified (**Table 4**). In total, 53% (45/85) of baits were removed from bait stations over 85 BENs, with 34 baits confirmed as ‘taken’ and 11 baits classed as ‘suspected taken’. Seventy-six percent (34/45) of the bait removals occurred on the first night they were placed in the field (**Figure 4**). Amongst identifiable animals, brushtail possums were responsible for the 20% (9/45) of the confirmed/suspected bait removals. Pademelons and eastern quolls were each responsible for 18% (8/45) of the confirmed or suspected bait removals. Devils were confirmed or suspected to remove 7% (3/45) of the removed baits.

**Figure 4.**
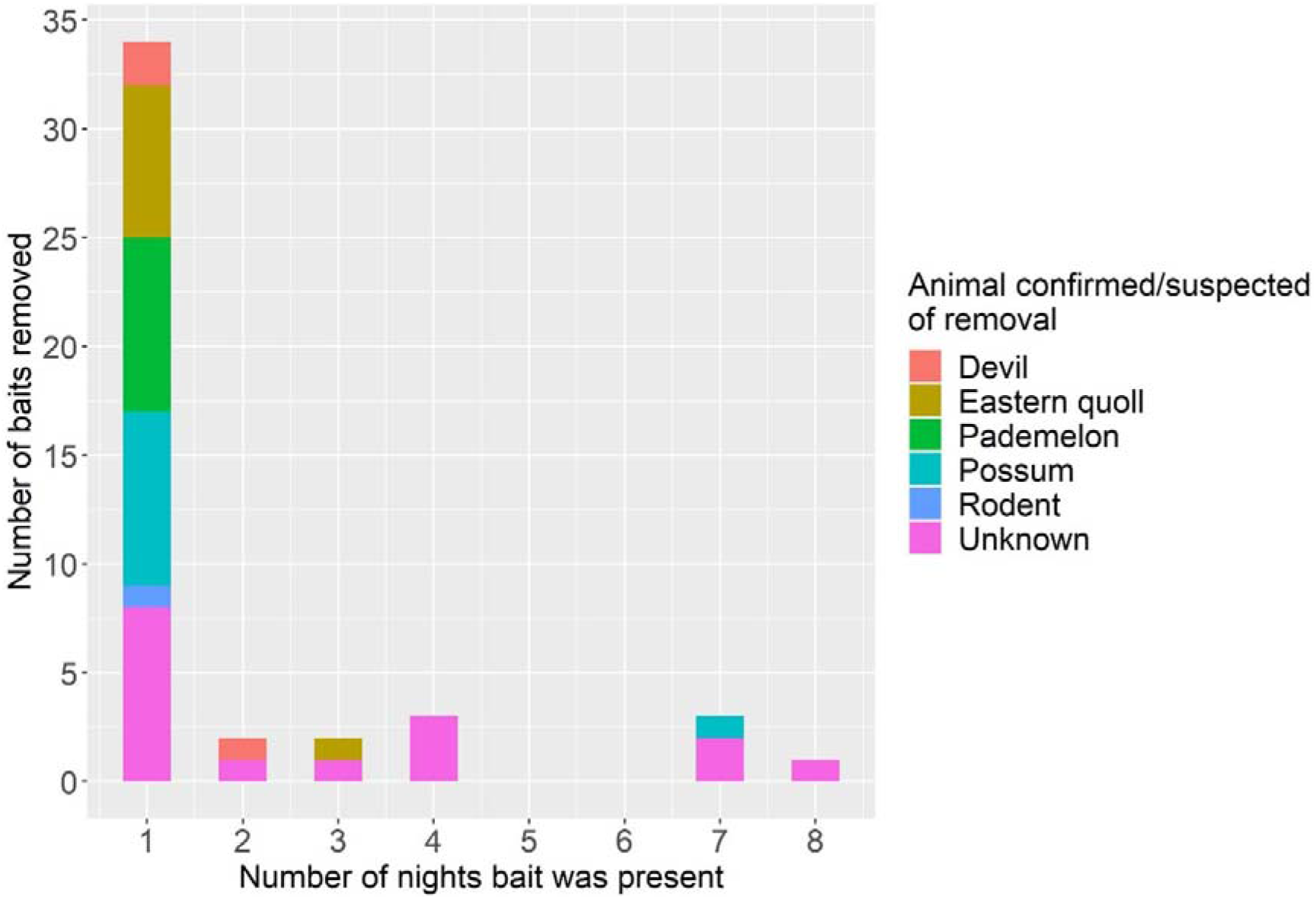
Consumption pressure by wild animals. Number of placebo baits confirmed or suspected to have been removed from bait stations over 85 bait exposure nights (BENs) across five baited study sites in southern Tasmania, 2021. 75% (34/45) of baits were removed on the first night. Confirmed and suspected removals by pademelons, brushtail possums, and eastern quolls accounted for a combined 55% (25/45) of the total baits removed from the bait station.

Bait stations with cereal baits were visited most frequently (5.4 visits/night) and comprised the largest number of devil visits (12) out of all four bait flavours. The cereal bait also had the most confirmed bait removals (11), including the only confirmed devil bait removal. Devils were also suspected to have removed one cereal and one fish bait, but no ruminant or kangaroo meat baits. Ruminant bait stations were the second most visited bait station (3.9 visits/night), with the second most baits confirmed taken (10) and three baits suspected taken.

Ruminant and kangaroo meat bait stations had the fewest visits by devils. Fish bait and kangaroo meat stations had 3 and 1.9 visits per night, respectively. Nine fish baits were confirmed taken from these bait stations, with four baits suspected taken (13 suspected/confirmed).

### Dispenser trials

As a result of the higher density non-target species taking most baits before lower density devils visited the bait stations, automated baited dispensers were evaluated at three of our local field sites. Automated bait dispenser were visited 270 times by animals from 10 categories over 33 BENs across three sites (**Table 5)**. Macropods were again the most frequent visitors (94), followed by Tasmanian devils (41), eastern quolls (41), rodents (41), brushtail possums (29), and cats (14). Rabbits, dogs, other marsupialsand unidentifiable animals were infrequent visitors to dispenser sites. A total of 112 dispenser interactions were observed by animals from five categories. Eastern quolls were responsible for the most dispenser interactions (39), followed by rodents (22), Tasmanian devils (21), macropods (20), brushtail possums (5) and cats (5). Devils accounted for 8% (21/270) of visits to bait dispensers compared to 6% (20/325) of visits to ground bait stations in the prior trials. However, devils removed 52% (11/21) of the baits from the dispensers (**Figure 5**) compared to only 6% of the ground baits. The percentage of baits retrieved by eastern quolls was 38% (8/21) from dispensers compared to 21% of ground baits. The bait dispensers decreased the percentage of baits retrieved by possums and no macropods retrieved a bait from the dispenser despite being the most common visitors in the dispenser videos.

**Table 5.** Dispenser interactions. Total amounts of visits, dispenser interactions, bait retrieval and suspected bait retrieval by animals/categories of animals from 33 bait exposure nights (BENs) across 3 sites

| Species or group | Visits | Dispenser interactions | Bait retrievals | Suspected retrievals |
| --- | --- | --- | --- | --- |
| Macropods | 94 | 20 | 0 | 0 |
| Brushtail possum | 29 | 5 | 1 | 0 |
| Eastern quoll | 41 | 39 | 8 | 0 |
| No ID | 9 | 0 | 0 | 0 |
| Tasmanian devil | 33 | 21 | 11 | 2 |
| Other marsupials | 4 | 0 | 0 | 0 |
| Rabbit | 3 | 0 | 0 | 0 |
| Cat | 14 | 5 | 0 | 0 |
| Dog | 2 | 0 | 0 | 0 |
| Rodents | 41 | 22 | 1 | 1 |
| Total | 270 | 112 | 21 | 3 |

**Figure 5.**
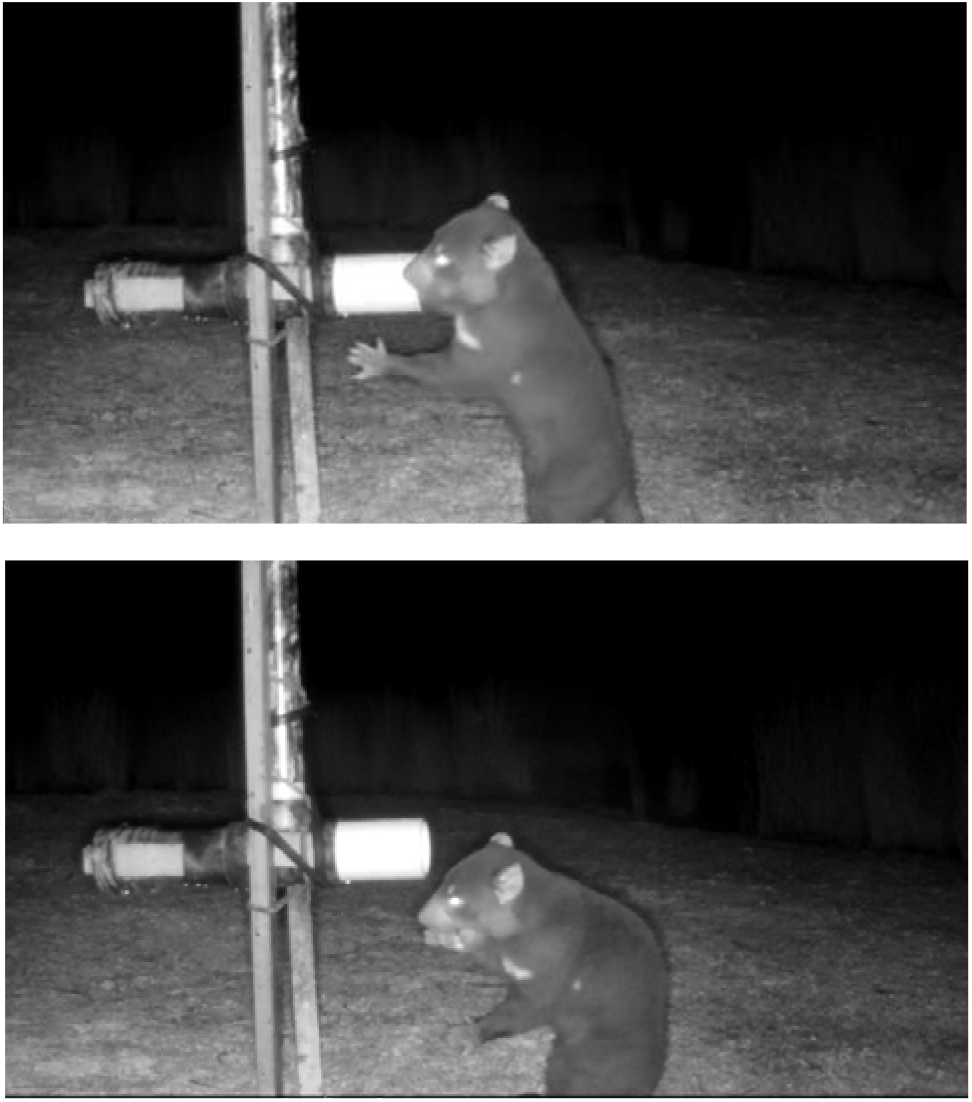
Devil retrieving bait from dispenser. Images from trail camera videos showing a Tasmanian devil retrieving bait from an automated bait dispenser and eating the retrieved bait. Links for the full videos are available in the Supplemental Material.

## Discussion

An oral bait vaccination strategy has been proposed for combatting the disease-induced decline in the wild devil population (Flies et al., 2020). However, very little information is available to assess the feasibility of this strategy from a vaccine bait delivery viewpoint (Hughes *et al*. 2011; Mallick *et al*. 2016). This study showed that manufactured baits consisting primarily of ruminant or fish meat were palatable to both captive and wild devils. However, bait modifications or automated dispensers are needed to reduce consumption by non-target species and improve specificity for devils.

In contrast to previous research that reported devils were overwhelmingly the most frequent visitors to ruminant-based bait stations and consumed the majority of baits in the field (Hughes *et al*. 2011), macropods and brushtail possums often arrived first and consumed the most baits in our study. We hypothesise that this difference is due in part to differences in species abundance and ecology between the studies. The Hughes et al. (2011) study occurred in a more remote part of northern Tasmania where devil density was high prior to devil facial tumour disease arriving in the area. Local devil populations have declined by approximately 80% across most of the state since that time (Lazenby *et al*. 2018; Cunningham *et al*. 2021). Most ground baits were removed on the first night, with more abundant species like pademelons and possums removing the baits before devils had an opportunity to locate and consume the bait.

We emphasize that field sites used in this pilot study were peri-urban and additional testing in other habitat types will be needed to develop effective regional baiting strategies. Future studies should occur at sites where devil and competitor populations densities have been quantified. Additionally, baiting near landscape and habitat features preferred by devils could also help improve uptake of ground baits by devils. For example, devils often travel along fence lines, whereas quolls are less likely to follow fence lines (Andersen *et al*. 2017).

The reduced devil density has been associated with other ecological changes, such as reduced fear behaviour in possums (Cunningham *et al*. 2019), which could also be associated with increase uptake by possums and other non-target species. Lower devil density may also coincide with an overabundance of devil food in the environment (e.g. macropods and possums as roadkill). Overabundance of devil food may lead to reduced devil foraging and interest in small food items like baits. Future studies at larger scales should be performed at long-term monitoring sites to assess the effects of devil density on the proportion of baits retrieved by devils.

Our use of automated bait dispensers increased the percentage of baits consumed by devils nearly ten-fold in this pilot study. Importantly, the design of the dispenser decreased bait consumption by macropods from 25% to 0%. This preliminary study also suggests that modifications to the dispenser position, such as raising the bait receptable higher off the ground could reduce uptake by small species. For example, we observed one wild rat retrieve a bait at 200 mm above the ground, but what appeared to be the same rat was unable to retrieve a bait once the receptacle of the dispenser had been raised to 350 mm. Eastern quolls were able to retrieve baits from a height of 350 mm but this often required a lunge to reach the bait, whereas devils could reach in and grab the bait at 350 mm. Future studies should test additional dispenser heights and refinement to the receptacle designs and non-target filters to maximize bait uptake by devils.

Dispensers suffer from the limitation that some individuals could repeatedly visit the dispensers throughout the night. We were unable to definitively identify individual devils in the dispenser footage, but it is very likely that some individual devils and quolls were able to retrieve more than one bait per night. Discriminatory sensor camera technology, as used in Felixer grooming traps (Read *et al*. 2019), may be used in conjunction with dispensers so that baits are only released when activation sensors recognise the shape of a devil. However, it seems unlikely that that image recognition technology could effectively identity individual devils to limit consumption of multiple baits by a single devil. Alternatively, topical chemical deterrents could be used with baits to limit the ability of individual animals to monopolise baits or food resources (Johnson *et al*. in press).

During wild trials a cat was observed at a dispenser site carrying a prey animal in its mouth. If future research shows dispenser sites attract a large numbers of prey animals, then predation by invasive predators could be an issue that would need to be addressed. For example, dispensers may be set up so for a brief period of time at a location.

Increasing target uptake could potentially be accomplished through habitat-based and seasonal baiting programs based on devil behavioural and movement ecology, as has been simulated for racoons (McClure *et al*. 2022). Likewise, non-target uptake by herbivores and omnivores could potentially be reduced by baiting during periods when other food sources are readily available. Current knowledge about spatial and temporal distributions of non-target species like pademelons (le Mar, 2002), brushtail possums (le Mar, 2002, le Mar & McArthur, 2005, Hollings et al., 2015), and eastern quolls (Hollings et al., 2013, Fancourt, 2010) would assist in timing bait distribution to correspond with periods when there is a greater ratio of target to non-target species.

Similar to a 2011 study that tested placebo ruminant-meat baits with captive devils (Hughes *et al*. 2011), we observed that male devils were more likely to consume baits than female devils at two captive devil facilities. A recent study suggested that males have larger home range sizes than females in areas affected by DFT1 (Comte *et al*. 2020), which could further lead to disproportionate uptake of baits by male devils. Studies of devil contact networks have shown that males are more likely to receive bite wounds than female (Hamilton *et al*. 2019). The wounds create the potential for pathogen transmission and are more commonly inflicted during the mating season when males mate-guard females. High uptake of an effective vaccine by males could prevent transmitted tumour cells from establishing in new male hosts or progressing towards more severe disease post-infection. Previous research has also shown that males with severe disease become increasingly socially isolated, likely due to the debilitating effects of the disease. Healthy, vaccinated male devils would be expected to be more competitive for mates and could ultimately reduce disease prevalence in females by reducing contact with diseased males.

Maximising delivery of the vaccine component of the bait is also critical for an effective oral bait vaccine. For example, if the vaccine component can be easily separated from the bait matrix, then ingestion of the vaccine will be reduced. The results of capsule trials show that most captive devils will eat and fully consume capsule-loaded fish bait. However, in many trials the capsule became separated from the bait matrix, which led to the bait being consumed but not the capsule. A uniform distribution of the vaccine throughout the bait matrix would force vaccine ingestion with the consumption of the bait matrix.

Feeding behaviour of the target species is also important for an effective vaccine. A study observing bait flavour preferences of captive raccoons and skunks with placebo baits found that raccoons were more likely to ‘chew the entire bait and hold the bait in their paws when eating’ than skunks (Johnson et al., 2016). Comparatively, skunks were observed to pin the bait on the ground and possibly chew the bait more than raccoons, increasing the chance of liquid vaccine dispersing onto the ground. In agreement with a prior prey-processing study of captive devils (Pollock et al., 2021), we found that captive and wild devils were likely to hold the bait in their paws and crush the bait.

In summary, the outcomes of this study lay the foundation for an oral bait vaccine delivery system if an effective DFT1/2 vaccine can be developed. A bait that is less attractive to non-target species will be needed before large scale ground bait campaigns could be implemented due to the cost of oral bait vaccines. Flavour preference and feeding behaviour information may be used to optimise bait matrix formulations by designing a palatable bait that cannot be easily consumed by non-target species. However, our study also suggests that bait dispensers could be used effectively at regional levels as a means of reducing non-target uptake and increasing the specificity of delivering currently available bait formulations to devils.

## Acknowledgments

We wish to thank Ginny Ralph for ongoing care of captive Tasmanian devils. We thank Androo Kelly and the knowledgeable keepers at the Trowunna Wildlife Sanctuary. We thank the Dave Schaap and Drew Lee and other members of the Tasmanian Government’s Save the Tasmanian Devil Program for coordinating access to captive devils. We thank Dr Linton Staples for providing baits and a wealth of knowledge of baiting practices. We thank A/Prof Chris Burridge, Prof Menna Jones, Prof Greg Woods, and A/Prof Bruce Lyons for the comments and support on the honours thesis associated with this study. We thank Dr Marissa Parrott for providing the base ethogram and comments on methods. We thank Dr Barrett Wolfe for statistical advice. We thank Robb Meijers and Tim Long for assistance with field trials.

## Ethical approval

The research study was approved by the University of Tasmania Animal Ethics Committee (#23220) and the Department of Natural Resources and Environment Tasmania Save the Tasmanian Devil Program’s Captive Research Advisory Group (‘Bait preference testing in Tasmanian devils’).

## Author contributions

ASF, RJP, SBA, SD, and TJS designed the study. ASF, JMM, RP, and SD performed the experiments. ASF, NMFJ, and SD performed the data analysis. ASF, ATG, NMFJ, RJP, SD, and TJS interpreted the results. ASF and SD wrote the manuscript, and all authors edited the manuscript.

## Data availability

Data and videos related to this paper will be publicly available through the Supplementary Materials and the University of Tasmania Research Data Portal: https://rdp.utas.edu.au/ using data key “RD-EOBDMFTDH”.

