## Supplementary Materials 1 for "Evaluation of oral baits and distribution methods for Tasmanian devils (*Sarcophilus harrisii*)"

**Title**

**Affiliations**

**Correspondence**

**Supplementary Materials**

**Table S1.** Devil behaviour ethogram displaying behaviours observed, the type of behaviour (either a discrete point event or continuous state event), and how that behaviour was defined.

| | **Behaviour** | **Behaviour type** | **Description** |
|---|---|---|---|
| **Interaction behaviours** | Bait approached | Point | Devil approaches bait to within touching distance, but does not make contact |
| | Bait contacted | Point | Devil makes physical contact with the bait |
| | Enrichment | Point | Devil is interacting with, but not eating, the bait, e.g. pushing bait around with paws |
| | Bait interaction: obstructed view | State | Devil interacts with bait but has its back turned or view is obstructed from the camera |
| | Bait sniffed | State | Devil sniffs bait and either makes contact or rejects bait |
| **Eating behaviours** | Bait pinned to ground | State | Devil uses paws to hold bait on the ground and attempts to bite into bait matrix |
| | Bait held in paws | State | Devil uses paws to hold bait and attempts to bite into the bait matrix |
| | Bait capsule punctured | Point | Bait capsule is punctured by devil and solution spills out |
| | Bait nibbled | State | Devil nibbles or pulls bait using incisors |
| | Bait crushed | State | Devil attempts to crush bait with its molars |
| | Bait licked | State | Devil licks bait either on the ground or in its paws |
| **Grooming behaviours** | Lick paws | State | Devil licks one or both paws |
| | Scratch body | Point | Devil uses nails to scratch body |
| | Scratch mouth | Point | Devil uses nails to scratch the interior of their mouth |
| | Face wipe | State | Devil grooms face with either one or both front paws; may be preceded by animal licking its leg first |
| **Other** | Scent mark | Point | Devil drags cloaca over/around area where baits were placed |
| | Confrontation | Point | In a pen with multiple devils, a devil exhibits vocalising or mouthing in competition for bait |
| | Capsule contents consumed | State | Sucrose from capsule is consumed by devil |

**Table S2**. Percentages of each consumption group for each bait flavour in single-housed devils from four palatability trips. 'Removed' baits indicate a bait that was taken away from the scope of the camera and could not be found after the pen was inspected; removed baits were not include in statistical analysis.

| Bait flavour | Uneaten | Partially eaten | Completely consumed | Removed |
|---|---|---|---|---|
| **Ruminant** | 25.5% (12/46) | 17.0% (8/46) | 57.4% (27/46) | 7 |
| **Fish** | 34.2% (13/38) | 13.1 (5/38) | 52.6% (20/38) | 12 |
| **Kangaroo** | 0.0% (0/6) | 0.0% (0/6) | 100.0% (6/6) | 1 |
| **Cereal** | 52.5% (21/40) | 15.0% (6/40) | 32.5% (13/40) | 3 |

**Table S3**. Total visitation from six study sites and one control site in wild trials for all 289 nights; this includes nights when baits were both present and not present.

| | Site | | | | | | | |
|---|---|---|---|---|---|---|---|---|
| Species or group | Sandfly 1 | Sandfly 2 | Sandfly 3 (control) | Lenah Valley 1 | Lenah Valley 2 | Ridgeway | Middleton | Total |
| Devil | 14 | 31 | 6 | 2 | 0 | 8 | 6 | 67 |
| Possum | 8 | 5 | 3 | 7 | 86 | 38 | 9 | 156 |
| Tasmanian pademelon | 154 | 263 | 141 | 28 | 41 | 141 | 0 | 768 |
| Bennett's wallaby | 2 | 2 | 0 | 40 | 0 | 10 | 0 | 54 |
| Cat | 3 | 1 | 0 | 4 | 0 | 0 | 2 | 10 |
| Dog | 0 | 0 | 0 | 2 | 0 | 3 | 0 | 5 |
| Native hen | 0 | 0 | 0 | 10 | 0 | 6 | 0 | 16 |
| Fantail | 0 | 0 | 0 | 0 | 0 | 2 | 0 | 2 |
| Blackbird | 0 | 0 | 0 | 1 | 0 | 0 | 0 | 1 |
| Tawny frogmouth | 0 | 0 | 0 | 0 | 1 | 0 | 0 | 1 |
| Brown bandicoot | 18 | 7 | 2 | 0 | 0 | 0 | 0 | 27 |
| Eastern barred bandicoot | 2 | 12 | 0 | 0 | 0 | 9 | 0 | 23 |
| Tasmanian bettong | 9 | 2 | 0 | 0 | 0 | 0 | 0 | 11 |
| Eastern quoll | 34 | 59 | 6 | 0 | 0 | 0 | 0 | 99 |
| Mouse | 0 | 0 | 1 | 0 | 0 | 0 | 0 | 1 |
| Rat | 0 | 0 | 0 | 1 | 0 | 0 | 0 | 1 |
| Rakali | 1 | 0 | 0 | 0 | 0 | 0 | 0 | 1 |
| Rabbit | 0 | 9 | 1 | 0 | 0 | 9 | 0 | 19 |
| No ID | 29 | 30 | 19 | 6 | 9 | 6 | 0 | 99 |
| Total visits | 274 | 421 | 179 | 101 | 137 | 232 | 17 | 1361 |

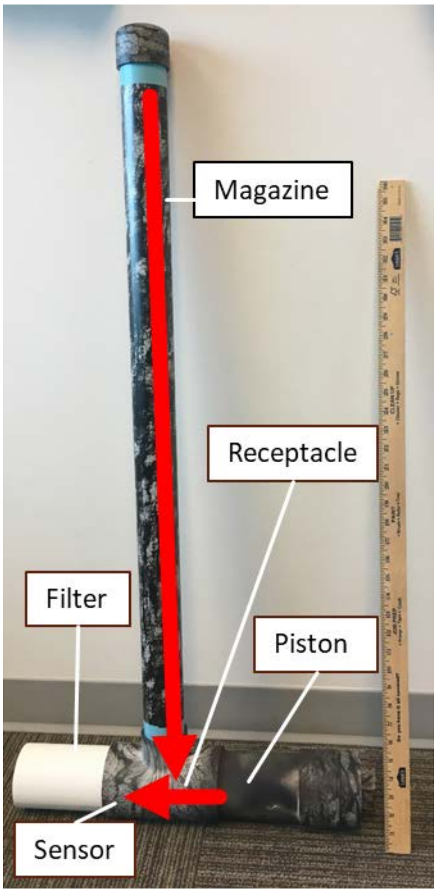


**Figure S1**. Automated bait dispenser used for wild dispenser trials. Developed by United States Department of Agriculture - National Wildlife Research Center. 1 meter ruler for scale.

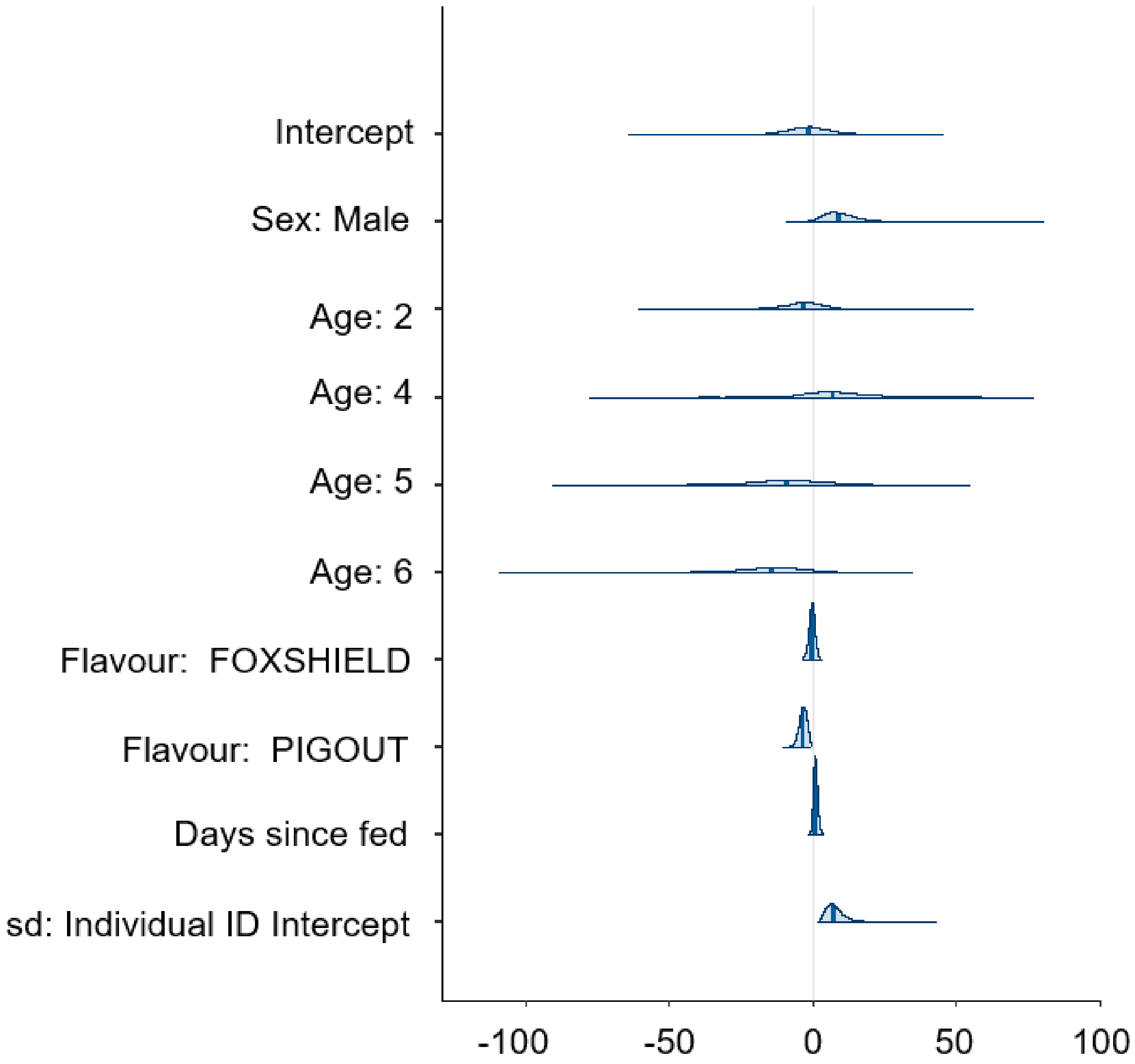


**Figure S2**. Log odds estimates of group-level and population-level effects of Model 1. Dark blue vertical lines represent the log odds estimate of a variable. The light blue shading indicates the limits of each variable’s 95% credible intervals.

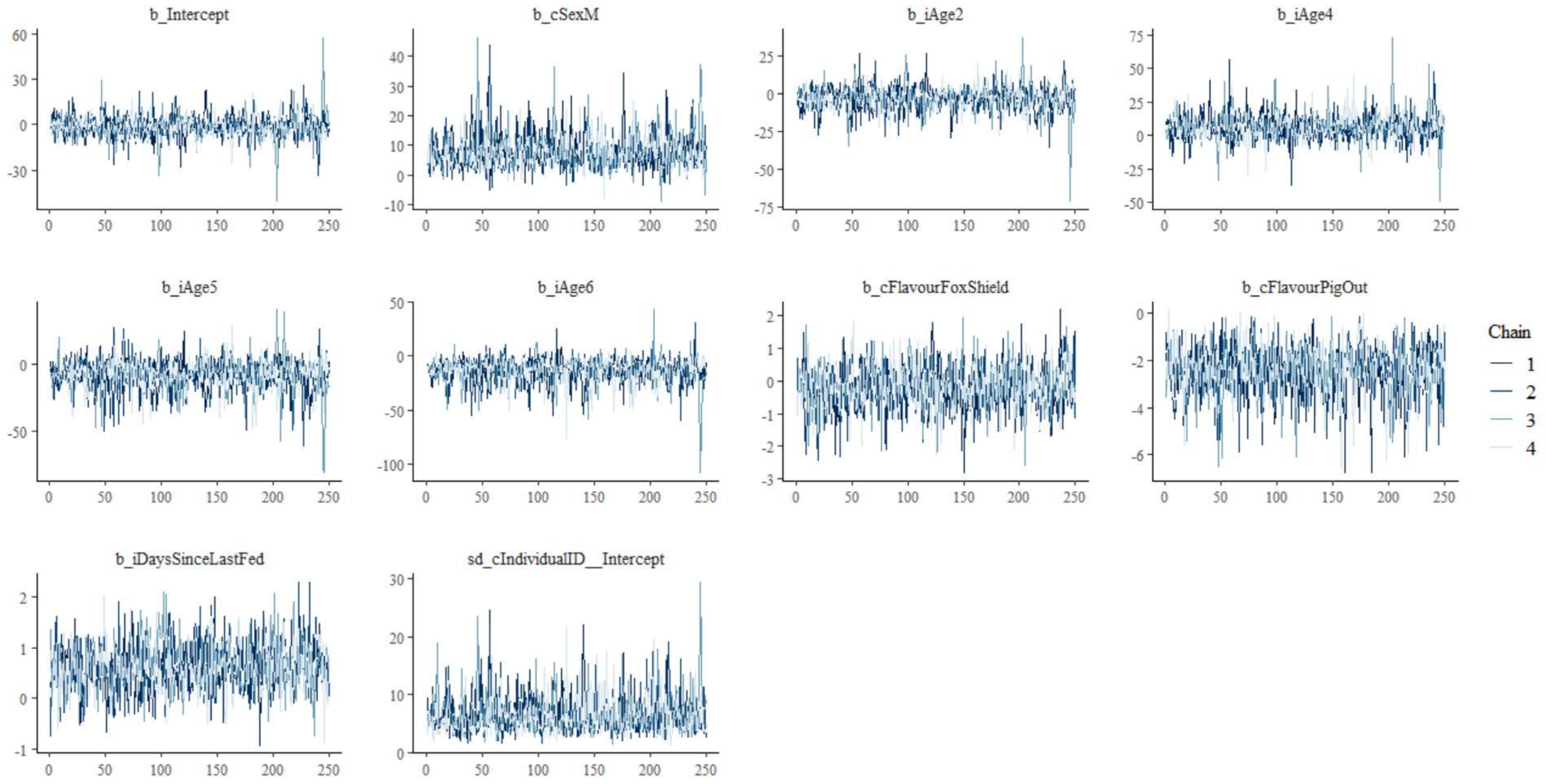


**Figure S3**. Convergence plot of Model 1. No divergences were evident in the models.

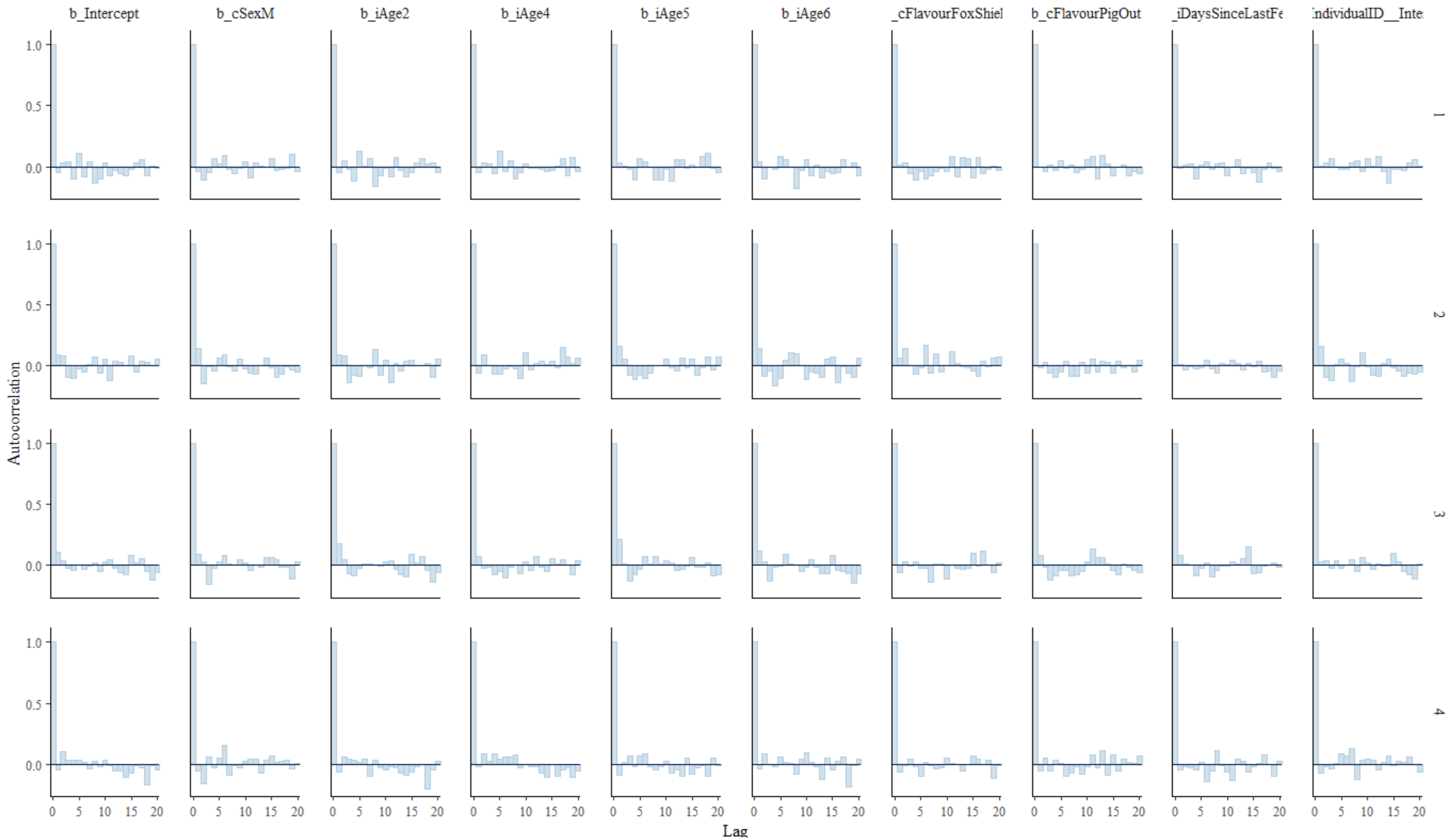


**Figure S4**. Autocorrelation plot of Model 1. No autocorrelation was evident in the models.

Supplemental video links:

Devil retrieving bait from dispenser (Sandfly)
https://youtu.be/5BEBfFqOY8k

Eastern quoll retrieving bait from dispenser (Sandfly)
https://youtu.be/4hBxDnZic8w
